# The *Drosophila* orthologue of the primary ciliary dyskinesia-associated gene, *DNAAF3*, is required for axonemal dynein assembly

**DOI:** 10.1101/2021.05.11.443597

**Authors:** Petra zur Lage, Zhiyan Xi, Jennifer Lennon, Iain Hunter, Wai Kit Chan, Alfonso Bolado Carrancio, Alex von Kriegsheim, Andrew P. Jarman

**Author notes:** These authors contributed equally to this work (Petra zur Lage, Zhiyan Xi). **Correspondence:** Andrew Jarman.

## Abstract

Ciliary motility is powered by a suite of highly conserved axoneme-specific dynein motor complexes. In humans the impairment of these motors through mutation results in the disease, Primary Ciliary Dyskinesia (PCD). Studies in *Drosophila* have helped to validate several PCD genes whose products are required for cytoplasmic pre-assembly of axonemal dynein motors. Here we report the characterisation of the *Drosophila* homologue of the less known assembly factor, *DNAAF3*. This gene, *CG17669* (*Dnaaf3*), is expressed exclusively in developing mechanosensory chordotonal (Ch) neurons and spermatocytes, the only two *Drosophila* cell types bearing motile cilia/flagella. Mutation of *Dnaaf3* results in larvae that are deaf and adults that are uncoordinated, indicating defective Ch neuron function. The mutant Ch neuron cilia of the antenna specifically lack dynein arms, while Ca imaging in larvae reveals a complete loss of Ch neuron response to vibration stimulus, confirming that mechanotransduction relies on ciliary dynein motors. Mutant males are infertile with immotile sperm whose flagella lack dynein arms and show axoneme disruption. Analysis of proteomic changes suggest a reduction in heavy chains of all axonemal dynein forms, consistent with an impairment of dynein pre-assembly.

**SUMMARY STATEMENT:** DNAAF3 function as a dynein assembly factor for motile cilia is conserved in *Drosophila*.

## INTRODUCTION

Motile cilia and flagella are highly conserved among eukaryotes ranging from unicellular organisms (e.g., *Chlamydomonas, Tetrahymena*) to mammals. A motile cilium or flagellum normally comprises a microtubule-based axoneme with nine peripheral microtubule doublets typically surrounding a central pair (9+2 structure). These two parts are connected by radial spokes. Some motile cilia lack the central pair and radial spokes (9+0 structure), such as the cilia in the embryonic node. In both 9+2 and 9+0 motile cilia, the components responsible for motility are the rows of cilium-specific (axonemal) dynein motor complexes, visible ultrastructurally as outer and inner dynein arms (ODA and IDA). These complexes drive the sliding of adjacent microtubule doublets to generate cilium movement (Porter, 2017). Dynein motors are large multisubunit complexes comprising heavy chains (HC, > 400 kDa) for force generation through ATP hydrolysis, intermediate chains (IC, 45-110 kDa) that scaffold the complex, and light chains (LC, < 30 kDa) that regulate motor activity (King, 2016).

The autosomal recessive genetic disease, Primary Ciliary Dyskinesia (PCD; MIM 244400), has seemingly complex clinical manifestations including chronic lung infections, progressive damage to the respiratory system, impaired male fertility and abnormal organ symmetry. In recent years the genetic basis of PCD has been intensely studied. The underlying cellular defect of PCD is the impaired motility of motile cilia/flagella, but this can result from mutation of one at least 50 different genes (Mitchison and Valente, 2017). Mutations of genes for certain axonemal proteins cause PCD, including several HC and IC subunits of the beat-generating ODA complex. Of interest are PCD mutations that identify genes required not for dynein components, but for dynein complex assembly and transport. Three stages are thought to constitute the biogenesis of dynein arms: cytoplasmic pre-assembly of multiple subunits into dynein complexes (Fok et al., 1994; Fowkes and Mitchell, 1998), transfer of the complexes into the ciliary compartment, and intraflagellar transport (IFT) along the axoneme (Horani and Ferkol, 2013). The pre-assembly of the dynein HCs may be further divided into two phases: the folding and stabilisation of globular head domains of HCs, and the assembly of HCs onto the IC dimer scaffold with incorporation of LCs (Mitchison et al., 2012). A distinct set of around ten proteins, categorized as axonemal dynein assembly factors (DNAAFs), is required for these steps. Their mutations typically cause a combined failure of outer and inner dynein arm localization on the axonemal microtubules, and the resultant loss of dynein motors can be observed by transmission electron microscopy (TEM) in the cross sections of respiratory epithelial cilia from mutation-bearing PCD patients (Mitchison and Valente, 2017).

Several DNAAF proteins contain domains that are found in HSP90 co-chaperones, such as PIH (Protein Interacting with HSP90), TPR (Tetratricopeptide Repeat), and CS (CHORD-SGT1), leading to the hypothesis that they mediate interactions with chaperones in the cytoplasm for the correct folding/assembly of axonemal dyneins. Several of these assembly factors are hypothesized to form co-chaperones similar to the well-known R2TP complex (Kakihara and Houry, 2012; Vaughan, 2014). Such putative ‘R2TP-like’ complexes may be formed by the PCD proteins DYX1C1 (DNAAF4) and SPAG1 (Knowles et al., 2013; Tarkar et al., 2013) with DNAAF2 (KTU) (Omran et al., 2008) and PIH1D3 (Paff et al., 2017; Tarkar et al., 2013). In *Drosophila*, WDR92 has been recently linked to R2TP function during dynein assembly (zur Lage et al., 2018). Separately, the PCD protein ZMYND10 is involved in HC stabilization during which it recruits co-chaperone FKBP8 (Mali et al., 2018). Overall, the molecular evidence supports the notion that many DNAAFs function as cytoplasmic cochaperones.

However, the interactions and functions of several assembly factors are not clear. DNAAF3 is such a factor. The human protein has no characteristic chaperone-related domains; the functions of its domains (DUF4470 and DUF4471) are not known. Loss-of-function mutations in *DNAAF3* were identified initially in three PCD cases that mapped to locus *CILD2* (MIM606763), which were characterised by an absence of outer and inner dynein arms in respiratory cilia and consequent ciliary immotility (Mitchison et al., 2012). In *Chlamydomonas* the DNAAF3 orthologue, PF22, is also required for the presence of ODAs/IDAs in flagella (Mitchison et al., 2012). Based on PF22’s cytoplasmic location, and protease sensitivity and antigen exposure of HCs in mutant cells, it was hypothesised that abnormal dynein complexes assemble and accumulate in *pf22* mutants, suggesting PF22 functions at a late step in dynein complex assembly, possibly in stabilising HCs or in late stage maturation after cytoplasmic pre-assembly (Mitchison et al., 2012). Apart from further human case reports (Guo et al., 2019), very little further has been published on *DNAAF3* or homologues.

*Drosophila* has recently emerged as a useful metazoan model of ciliary motility. Dynein motors and other ciliary motility components are highly conserved in *Drosophila* despite the fact that it has only two cell types bearing motile cilia/flagella: the sensory cilium of mechanosensory chordotonal (Ch) neurons and sperm flagellum (zur Lage et al., 2019). In both cell types, the motility machinery is critical for function: if defective, mutant flies are deaf and uncoordinated as the motors are required to generate force during Ch neuron mechanotransduction, and the males are infertile due to immotile sperm (zur Lage et al., 2019). As such, *Drosophila* is a useful model for identification and analysis of ciliary motility genes, including dynein assembly factors (Diggle et al., 2014; Moore et al., 2013; zur Lage et al., 2018). A single orthologue of *DNAAF3* exists in *Drosophila: CG17669* (hereafter referred to as *Dnaaf3*). Here we show that *Dnaaf3* is functionally conserved. Tagged *Dnaaf3* protein is confined to the cytoplasmic compartment of developing Ch neurons and spermatocytes. RNAi depletion and CRISPR-generated null alleles specifically exhibit phenotypes consistent with ciliary/flagellar immotility with loss of both ODA and IDA. Proteomic analysis of mutants confirmed a specific reduction in abundance of motor proteins, particularly HCs. Unlike reported for *Chlamydomonas*, it appears that all axonemal dynein types may be affected.

## RESULTS

### *CG17669* is a *DNAAF3* orthologue and is expressed exclusively in cells bearing motile cilia/flagella

The *Drosophila* genome has a single homologue of *DNAAF3. CG17669* encodes a predicted protein having 27% amino acid sequence identity and 43% similarity to human DNAAF3 (Fig. 1A). The *CG17669* protein retains the DUF4470 and DUF4471 domains present in human and *C. reinhardtii* proteins. Therefore, we name the gene *Dnaaf3*.

**Fig. 1.**
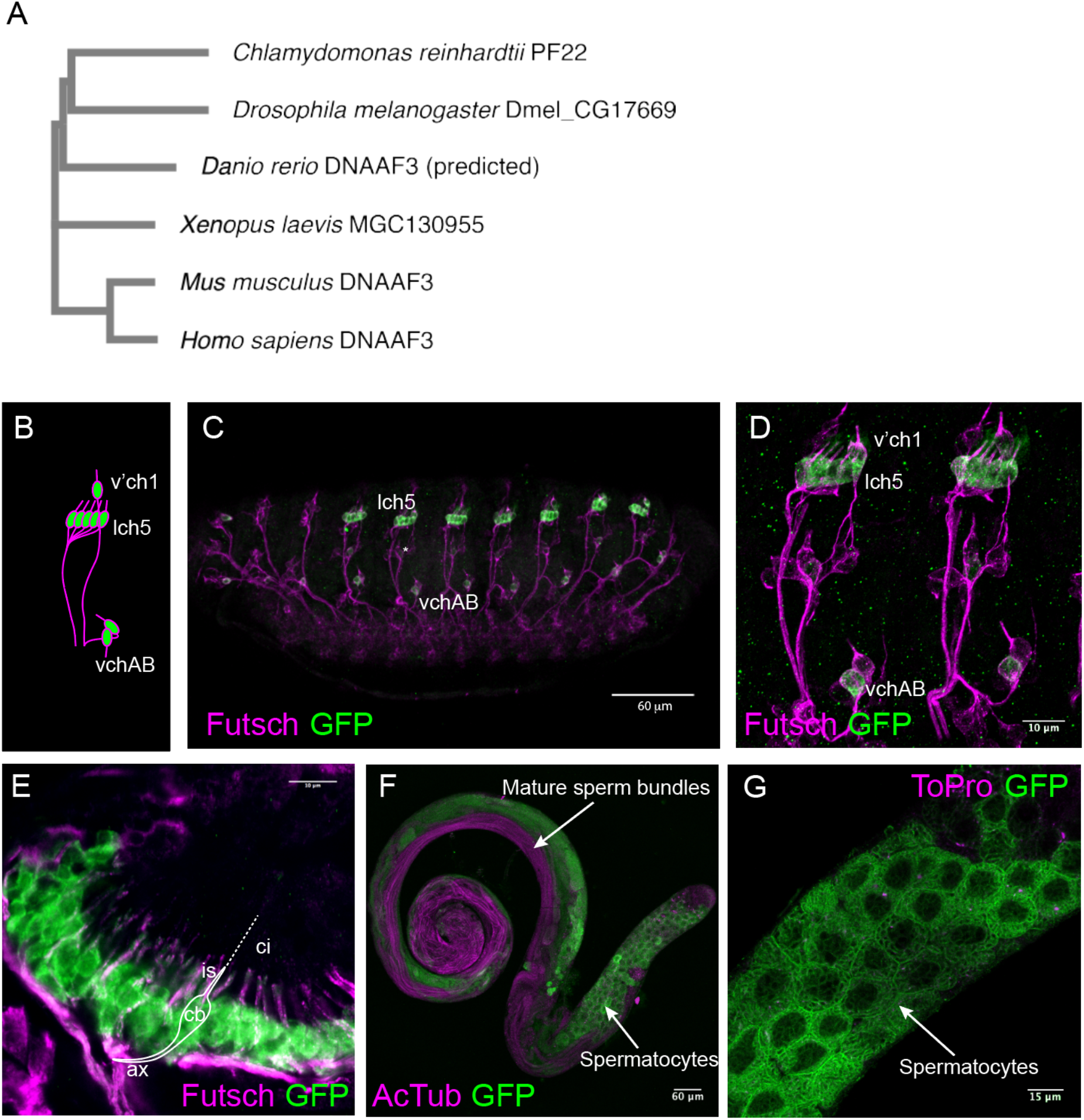
Expression of CG17669/Dnaaf3. (A) Phylogenetic relationship of *CG17669* with other *DNAAF3* orthologues, including PF22 from *Chlamydomonas*. Accession numbers for the proteins are: AEC04845 (for *C. reinhardtii* PF22), NP_611336 (for *Drosophila melanogaster* CG17669), XP_003201392 (for *Danio rerio* predicted DNAAF3), NP_001089839 (for *Xenopus laevis* MGC130955), NP_001028720 (for *Mus musculus* DNAAF3), NP_001243643 (for *Homo sapiens* DNAAF3, isoform 1). (B–G) Expression of Dnaaf3-mVenus fusion gene. (B) Schematic representation of Ch neuron arrangement in an embryonic abdominal segment, with clusters labelled. (C) Stage 16 embryo, showing expression in Ch neurons (anti-GFP, green) relative to sensory neurons (anti-Futsch, magenta). Scale bar: 60 μm. (D) Higher magnification view of two abdominal segments. Scale bar: 10 μm. (E) Pupal antenna, showing expression in the Ch neurons of Johnston’s organ. Expression is observed in the neuronal cell bodies (cb) and dendritic inner segment (is), but not the cilia (ci, unlabelled by Futsch). Axons (ax) are also indicated. Scale bar: 10 μm. (F) Adult testis showing fusion gene expression (green) in spermatocytes but not in mature sperm bundles (labelled with anti-acetylated tubulin, magenta). Scale bar: 60 μm. (G) Higher magnification view of testis showing reticulate pattern of fusion protein in spermatocytes (counter-label for nuclei is To-Pro-3, magenta). Scale bar: 15 μm.

To characterise *Dnaaf3* expression, we constructed a fly line that expresses a *Dnaaf3-mVenus* fusion gene, with mVenus in frame with the C-terminus of *Dnaaf3* protein (Fig. S1). The construct was designed to include the predicted tissue-specific promoter sequences of *Dnaaf3*, containing predicted binding sites for Rfx and Fd3F motile cilia transcription factors (consensus binding sequence RYYRYYN(1–3)RRNRAC and RYMAAYA respectively) (Laurençon et al., 2007; Newton et al., 2012). In the embryo, the fusion protein was detected in all differentiating Ch neurons (lch5, v’ch1, and vchAB), but not elsewhere, including other classes of sensory neurons that only have non-motile cilia (Fig. 1B–D). In Ch neurons, *Dnaaf3-mVenus* protein was localised to cytoplasm (including dendrites) but not the terminal cilia. Adult Ch neurons develop at the pupal stage during metamorphosis, notably within the developing antenna where they form the proprioceptive/auditory organ called Johnston’s organ. In immunofluorescence of pupal antennae, the fusion protein was expressed in the cytoplasm of Ch neurons exclusively (Fig. 1E).

In testes, the apical tip has spermatogonia that later undergo four mitotic divisions to form primary spermatocytes. The latter go through two meiotic divisions and produce spermatids. Spermatids undergo flagellogenesis within the cytoplasm and the flagellum is then extruded through a process of membrane remodelling and individualization to form mature sperm that are finally transferred to the seminal vesicle in a motility-dependent manner. The *Dnaaf3-mVenus* fusion protein was detected in spermatocytes and not in mature sperm (Fig. 1F). It appeared mostly in the cytoplasm in a reticulate pattern (Fig. 1G).

In summary, *Dnaaf3* protein is expressed in differentiating motile ciliated cells, and is located to the cytoplasm rather than the cilium/flagellum. The tissue pattern of *Dnaaf3* transcription was corroborated by *in situ* hybridisation: *Dnaaf3* mRNA is present specifically in embryonic differentiating Ch neurons and in spermatocytes of testes (Fig. S2).

### *Dnaaf3* is required for sperm motility

To address whether *Dnaaf3* is required for flagellar motility, we generated males with testis-specific *Dnaaf3* knockdown using the Gal4/UAS system (*Bam*-VP16-Gal4>UAS-*CG17669^GD36539^* RNAi). In a fertility assay, the knockdown males were observed to mate but produced no offspring (Fig. 2A). The testes of knock-down flies had a normal appearance with apparent sperm bundles (Fig. 2B–E). However, the seminal vesicles were devoid of motile sperm. Even upon crushing of testes, no motile sperm were released from mutant testes (Movies 1,2).

**Fig. 2.**
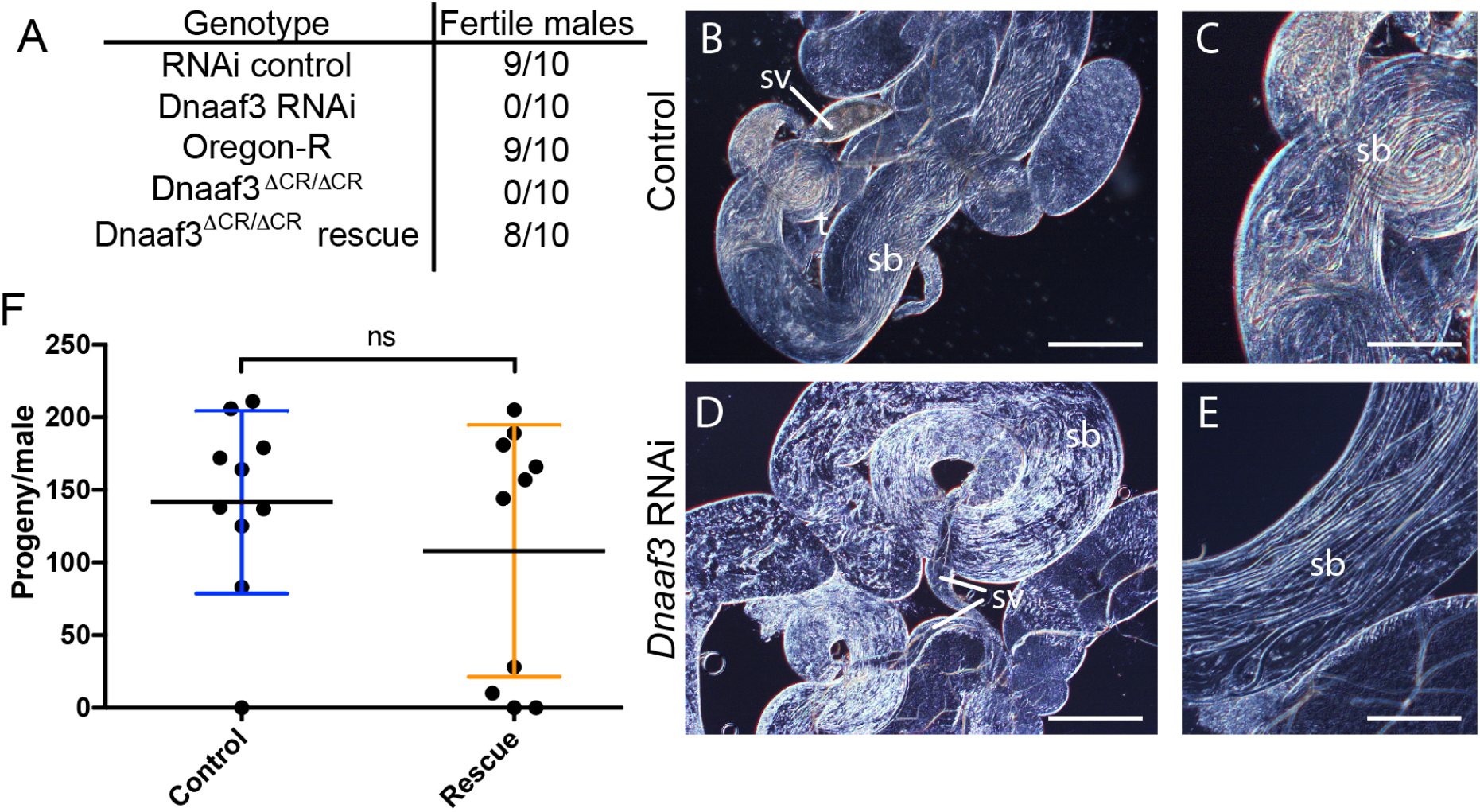
*Dnaaf3* is required for male fertility and sperm motility. (A) Male fertility, presented as number of tested males that produced offspring. (B–E) Adult testes examined by light microscopy. RNAi knockdown of *Dnaaf3* results in normal-looking testes with sperm bundles (sb) but no motile sperm are observed in seminal vesicles (sv). Scale bar: 50 μm (B,D) or 25 μm (C,E). (F) Fertility of *Dnaaf3^ΔCR^* homozygotes is substantially rescued by *Dnaaf3-mVenus* fusion gene. Graph shows number of progeny per male (n=10 males) for control and rescue (*Dnaaf3-mVenus/+; Dnaaf3^ΔCR^/Dnaaf3^ΔCR^*). The average progeny per male is not significantly different by two-tailed Mann-Whitney test (U=43.00; n=10; p=0.6299).

To confirm this phenotype, we generated a null allele of *Dnaaf3* by CRISPR/Cas9 catalysed gene replacement of the ORF with a *mini-white* gene. Analysis of homozygous *Dnaaf3^ΔCR^* mutants confirmed that males were viable but sterile, despite being able to mate (Fig. 2A). Their testes were normal in shape and contained normally elongating flagellar bundles of spermatids, but no motile sperms could be observed (Movie 3). Importantly, *Dnaaf3^ΔCR^* infertility could be substantially rescued by the *Dnaaf3-mVenus* fusion gene. Most rescued males (*Dnaaf3-mVenus/+; Dnaaf3^ΔCR^/Dnaaf3^ΔCR^*) produced progeny, with average fertility not significantly different from that of controls (Fig. 2F). Upon dissection, the seminal vesicles of the rescued males were observed to contain motile sperm (Movie 4).

### *Dnaaf3* is required for mechanotransduction by Ch neurons

The Ch neuron cilium is the site of mechanotransduction by these neurons, and this requires functional axonemal dynein motors (Karak et al., 2015; Newton et al., 2012). To assess whether the proprioceptive function of Ch neurons requires *Dnaaf3*, we generated flies with sensory-neuron-specific *Dnaaf3* knockdown (scaGal4>UAS-Dcr2/UAS-CG17669^GD36539^ RNAi). These knockdown flies were viable but showed uncoordinated locomotion, which was reflected in poor performance in a climbing assay (Fig. 3A). This is consistent with defective Ch neuron mechanotransduction. Homozygous *Dnaaf3^ΔCR^* mutant flies also showed uncoordinated locomotion in the climbing assay (Fig. 3B). This effect was rescued by the fusion gene (Fig. 3C).

**Fig. 3.**
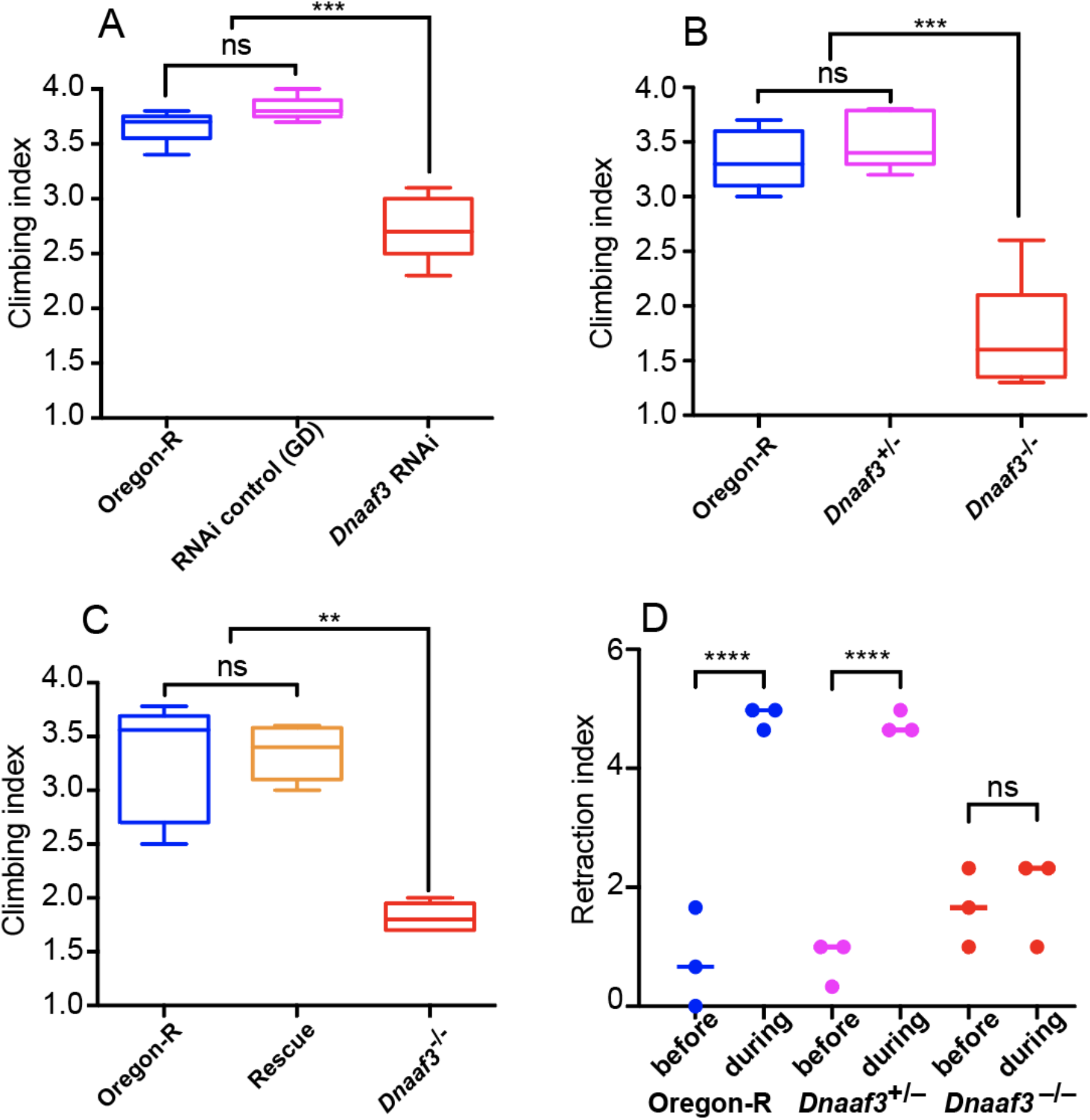
*Dnaaf3* is required for proprioceptive and auditory behaviours. (A–C) Climbing assays for testing adult incoordination as an indication of proprioceptive function of Ch neurons. N = 5 batches of 10 flies for each condition. Data are plotted as median and interquartile range. Significance was determined by Kruskal-Wallis test. (A) Knockdown of *Dnaaf3* in sensory neurons results in impaired climbing assay performance. *** indicates p = 0.0002. (B) *Dnaaf3^ΔCR^* homozygous null mutant flies show impaired climbing assay performance. *** indicates p = 0.0009 (C) The performance in the climbing assay of *Dnaaf3^ΔCR^* homozygous null mutant flies can be rescued by presence of Dnaaf3-mVenus fusion protein (‘rescue’). ** indicates p = 0.0027 (D) *Dnaaf3^ΔCR^* homozygous null mutant larvae are unresponsive to a 1000 Hz tone, consistent with defective Ch neurons. n = 3 batches of 5 larvae in each condition. **** indicates p<0.0001.

Ch neurons are also the receptors of auditory and vibration stimuli. For the larva, this function is reflected in its response to sound: a brief 1000-Hz tone causes larvae to momentarily retract, a response that depends on Ch neuron function (zur Lage et al., 2018) (Fig. 3D). In this assay, we observed that homozygous *Dnaaf3^ΔCR^* larvae do not respond to a 1000-Hz tone (Fig. 3D), suggesting failure in auditory/vibration mechanotransduction in Ch neurons.

To determine directly whether the larval Ch neurons can respond to a tone stimulus, we visualised Ch neuron activation in larval fillet preparations by recording calcium changes in their axon termini using a genetically supplied GCaMP calcium reporter (iavGal4, UAS-GCaMP6f). Freshly dissected 3^rd^ instar larval pelts were stimulated by 1024-Hz vibrations and the GCaMP response recorded by imaging. In *Dnaaf3^ΔCR^* heterozygote larvae, Ch neurons showed robust, short duration peaks in Ca-induced fluorescence upon vibration stimulus (mean peak response to stimulation, ΔF/F_0_: 13.54% ± 3.63; n = 14 stimuli in 5 larvae) (Fig. 4A,C,E). In contrast, Ch neurons from homozygous *Dnaaf3^ΔCR^* larvae did not show a robust response (mean peak ΔF/F_0_: 0.25% ± 0.22; *n* = 15 stimuli in 5 larvae) (Fig. 4B,D,F). There was a significant difference between mean peak ΔF/F_0_ in heterozygotes and homozygotes (unpaired t test, *P* ≤ 0.0001).

**Fig 4.**
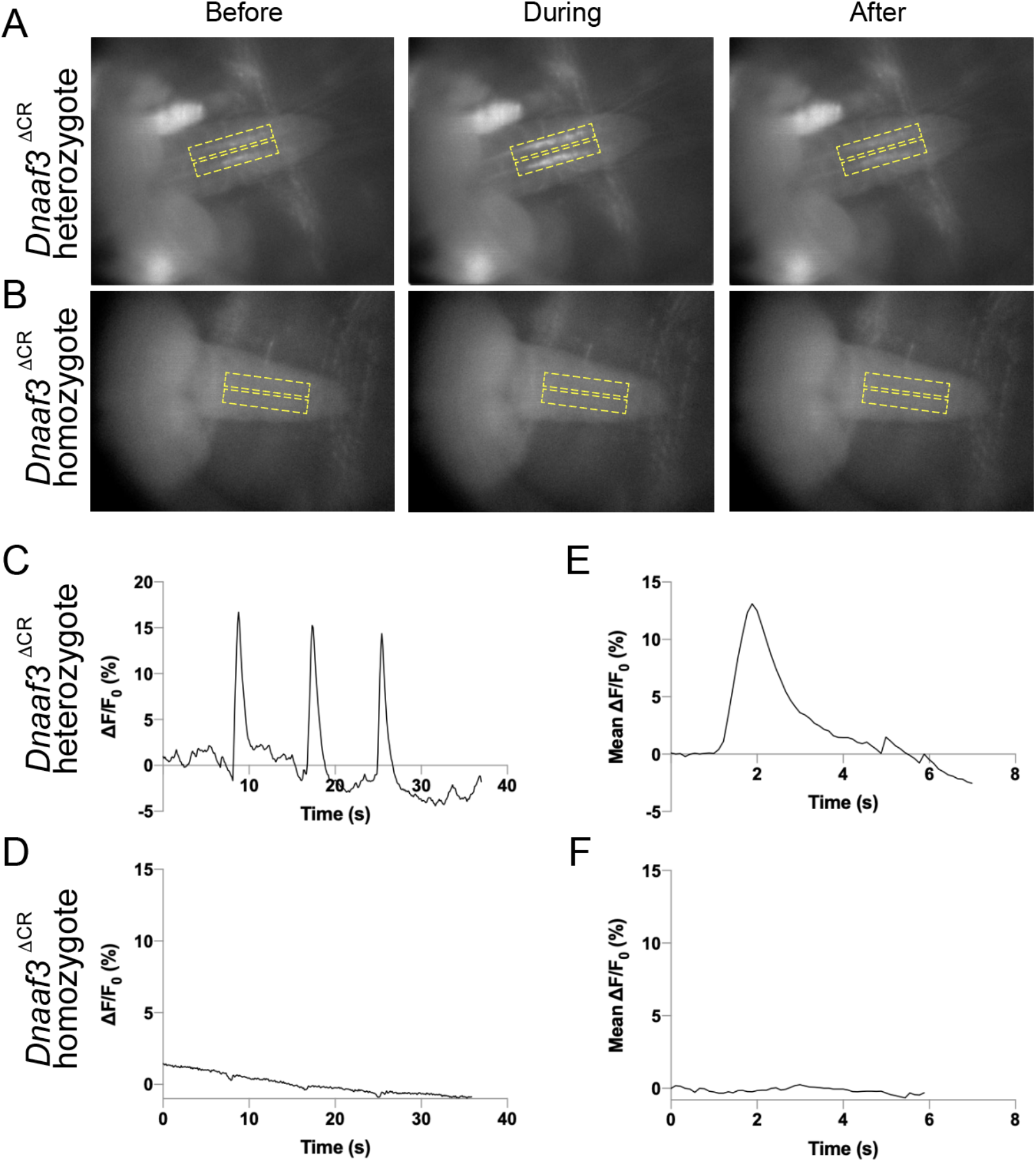
Response of larval Ch neurons to vibration stimulation is abolished in *Dnaaf3^ΔCR^* null mutants. Response to 1024Hz vibration stimulation, as fluorescence change in Ch axonal terminals in VNC of semi-intact larval preparations with Ch neuron-specific Gal4 driving GCaMP expression (iav-Gal4 x UAS-GCaMP6f). (A,B) Representative frames from video before, during and after stimulation for *Dnaaf3^ΔCR^* heterozygote (A) and *Dnaaf3^ΔCR^* homozygote (B). Dashed lines indicate region of interest defined to produce traces. (C,D) Representative traces for *Dnaaf3^ΔCR^* heterozygote (C) and homozygote (D) for response (ΔF/F_0_, %) to 3 successive 1-s stimuli. F_0_ is defined as mean F of: (0 s to onset of 1^st^ peak) + (end of 1^st^ peak to onset of 2^nd^ peak) + (end of 2^nd^ peak to onset of 3^rd^ peak). (E,F) Mean peak response for *Dnaaf3^ΔCR^* heterozygote (E, mean peak ΔF/F_0_ = 13.54% ± 3.63; n = 14 stimulations in 5 larvae) and *Dnaaf3^ΔCR^* homozygote peak (F, mean ΔF/F_0_ = 0.25% ± 0.22; n = 15 stimulations in 5 larvae).

Overall, null mutant individuals have multiple phenotypes consistent with impaired motility of cilia/flagella. This strongly supports the hypothesis that *Drosophila Dnaaf3* is required for dynein motor assembly or function.

### Ultrastructural analysis shows specific absence of dynein arms from cilia and flagella

Human *DNAAF3* mutations are characterised by ultrastructural absence of ciliary ODA/IDA, consistent with a failure in cytoplasmic pre-assembly of dynein motor complexes. We used TEM to analyse Ch neuron cilia and sperm flagella in *Dnaaf3^ΔCR^* mutant adult flies. In the antennal Johnston’s organ, transverse sections of Ch neuron cilia revealed a normal axonemal structure of 9+0 microtubule doublets, but ODA and IDA both appeared lacking (Fig. 5A,B). The loss of ODA was further examined in pupal Johnston’s organ Ch neurons by immunofluorescence using an antibody generated against the *Drosophila* orthologue of the ODA HC, Dnah5 (CG9492). Dnah5 protein was observed to be strongly reduced or absent in the cilia of Ch neurons in *Dnaaf3^ΔCR^* homozygote antennae (Fig. 6).

**Fig. 5.**
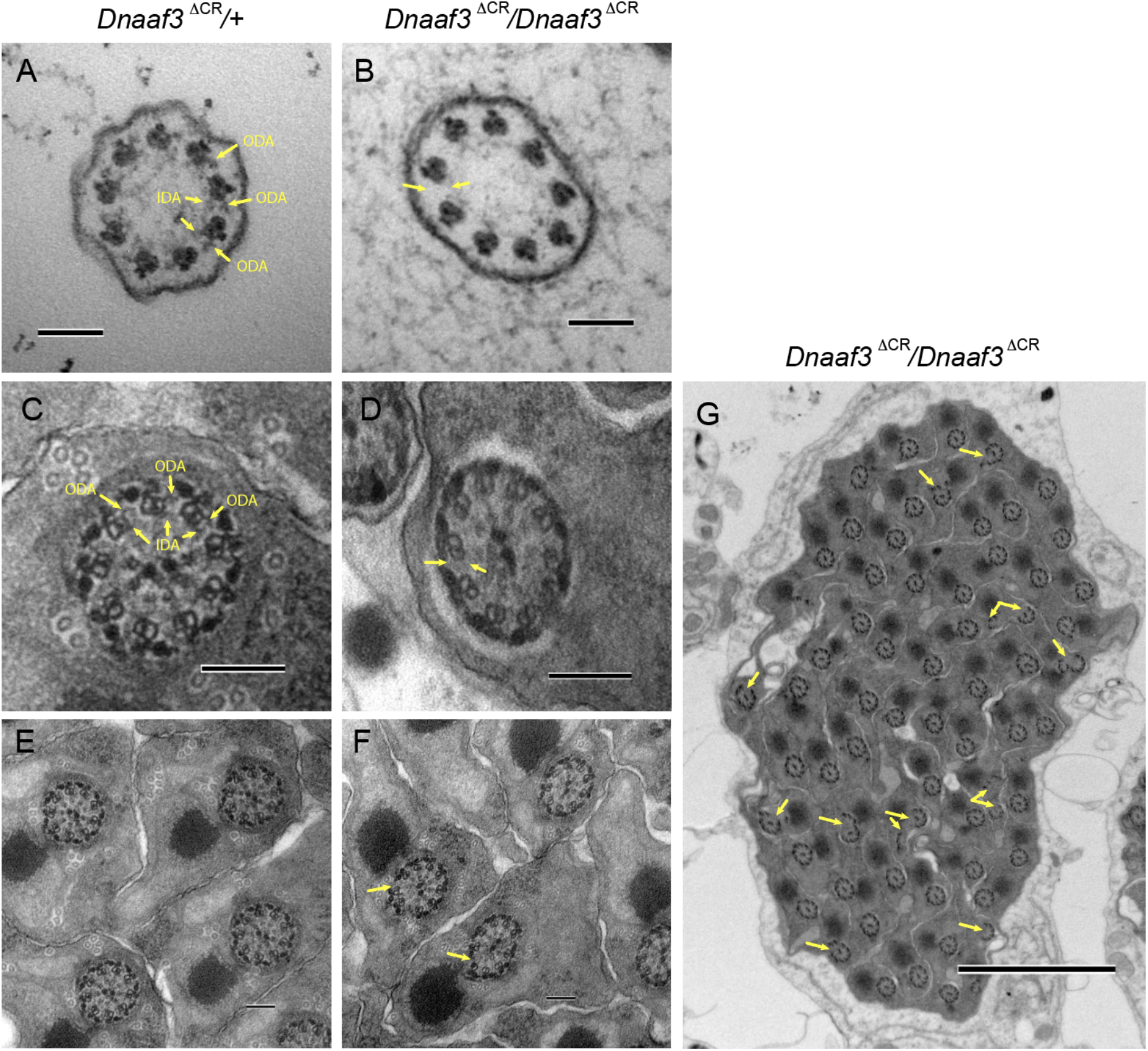
Loss of dynein arms in *Dnaaf3* mutant motile cilia and flagella. Transmission electron micrographs of adult antennal Ch neuron cilia (A,B) and testis sperm bundles (C–G), transverse sections. (A) *Dnaaf3^ΔCR^* heterozygote Ch neuron cilium, showing 9+0 microtubule doublets with attached outer and inner dynein arms (some arrowed, ODA, IDA). (B) *Dnaaf3^ΔCR^* homozygote Ch neuron cilium, microtubule doublets are intact, but lack visible ODA and IDA (example of expected location arrowed). (C) *Dnaaf3^ΔCR^* heterozygote sperm flagellum, showing 9+2 structure with ODA and IDA visible on at least some doublets. (D) *Dnaaf3^ΔCR^* homozygote sperm flagellum, showing regular axonemal structure but ODA/IDA are not visible. (E,F) Lower magnification views, with axonemal splits visible in homozygote (arrowed). (G) Low magnification transverse section of a mature sperm bundle from homozygote, showing generally normal bundle but with frequent axonemal splits (arrows). Scale bars: 100 nm (A–F); 2 μm (G)

**Fig 6.**
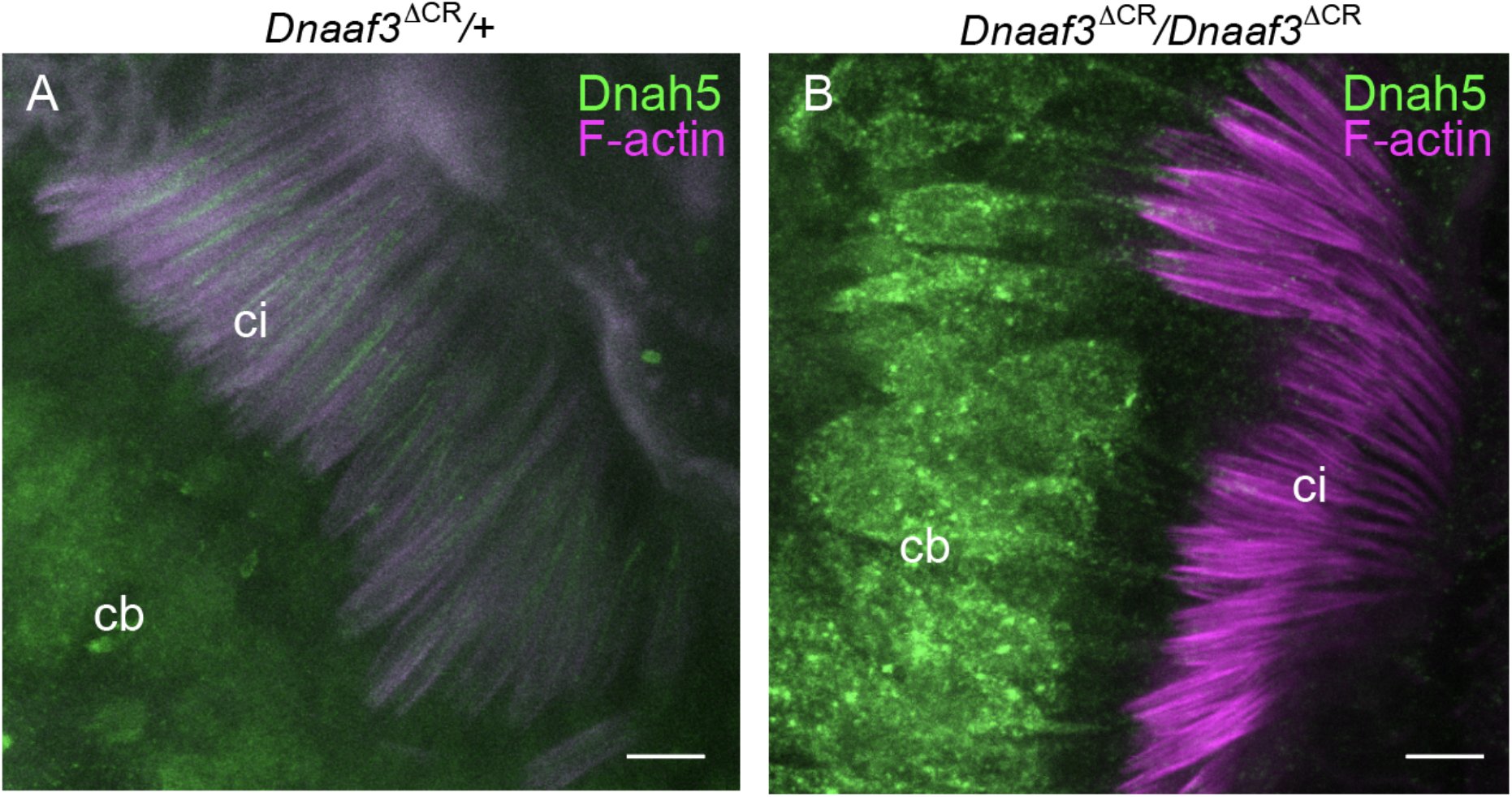
Loss of dynein heavy chain in *Dnaaf3* mutant motile cilia. Immunofluorescence imaging of chordotonal neurons in pupal antenna; Dnah5 protein (green) and phalloidin staining of F-actin (red), which marks the scolopale structures surrounding the ciliary dendrites. (A) *Dnaaf3^ΔCR^* heterozygote showing Dnah5 protein in cell bodies and cilia. (D) *Dnaaf3^ΔCR^* homozygote showing Dnah5 in cell bodies but absent from cilia. Number of antennae imaged: heterozygote n=15; homozygote n=10. Scale bars: 5 μm.

In the testes of *Dnaaf3^ΔCR^* homozygote males, TEM showed that sperm flagellum axonemes were largely normal, but strongly lacked ODA/IDA (Fig. 5C–F). In the sperm bundles, axonemal breakage was also observed, whereby one or more microtubule doublets break away from the axoneme (Fig, 5F,G). Such a phenotype has been observed for most other homologues of dynein assembly factors (zur Lage et al., 2018).

### Proteomic analysis of mutant testes reveals specific reduction in abundance of dynein heavy chains

In many dynein assembly factor mutants, failure in pre-assembly and localization of dynein motors appears to result in instability of some dynein subunits, particularly HCs (Mitchison et al., 2012). In *Drosophila*, testis protein abundances have previously been assayed by mass spectrometry in order to characterize the phenotype of the dynein assembly factor, *Wdr92* (zur Lage et al., 2018). We similarly carried out mass spectrometry on adult testes from *Dnaaf3^ΔCR^* mutant and control flies. By label-free quantitative MS, we detected 5549 proteins. To examine ciliary proteins, we focused on the MS data for homologues of proteins associated specifically with motile cilia (dynein motors, nexin-dynein regulatory complex, radial spokes, etc (zur Lage et al., 2019)). Of these 92 candidate proteins, 81 were detected as being present in the MS data but we excluded 17 that were not reliably detected (Table S1). Interestingly, in wild-type testes, dynein assembly factors and chaperones are among the most abundant proteins, suggesting that dynein assembly is a major activity in spermatocytes (Table S1).

Among the proteins detected is Dnaaf3 itself, and as expected this protein shows a large depletion in the *Dnaaf3^ΔCR^* mutant testes (Table 1). Of the ciliary motility proteins detected, 26 of them showed significant difference in abundance in *Dnaaf3^ΔCR^* mutant testes (>1.5-fold) (Table 1, Fig. 7A). Most of these were depleted, and notable among these are axonemal dynein HCs, of which all eight chains are significantly depleted (Table 2, Fig. 7B,C). A few ciliary proteins were enriched, and these are notably a subset of other dynein assembly factors (Wdr92, Dyx1c1, Lrrc6).

**Fig. 7.**
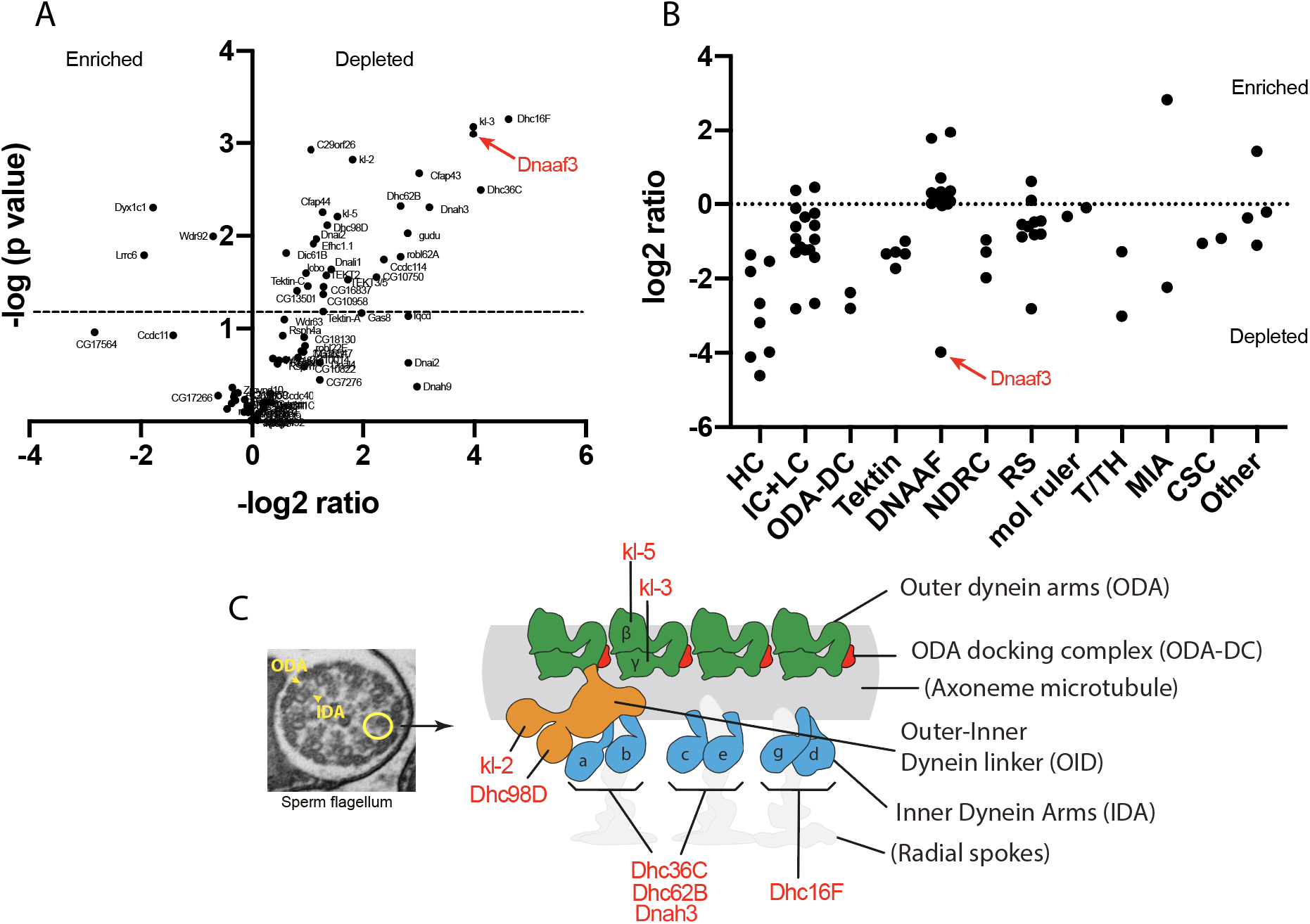
Dynein motor subunits are depleted in *Dnaaf3* mutant testes. (A) Volcano plot of motile cilia-associated proteins detected by MS in testes. To the left of the Y axis are proteins that are more abundant in homozygote testes (enriched); to the right are proteins that are less abundant (depleted). Dnaaf3 protein itself is strongly depleted as expected (red). (B) Summary of protein abundances for ciliary motility proteins. Below the 0 line are proteins depleted in the mutant. HC=dynein heavy chain; IC=dynein intermediate chain; LC=dynein light chain; ODA-DC= outer dynein arm docking complex; DNAAF=dynein assembly factor; NDRC=nexin-dynein regulatory complex; RS=radial spoke; mol ruler= molecular ruler complex (CCDC39/40); T/TH=tether/tether-head complex; MIA=modulator of inner arm; CSC=calmodulin-spoke complex. (C) Schematic of the arrangement of dynein motor complexes along one 96-nm repeating unit of an axonemal microtubule in the sperm flagellum (adapted from (Lin et al., 2014; zur Lage et al., 2019)). Colours indicate ODA (green), ODA-DC (red), double-headed IDA f (orange), single headed IDAs a–g (blue). The basal part of IDA f is also called the Outer-Inner Dynein linker (OID). The heavy chains of all subtypes (red protein names) are depleted in the mutant testes.

**Table 1.** Ciliary motility proteins whose abundance changes in *Dnaaf3* mutant testes (p<0.05), sorted by p value

| Majority protein IDs | Gene names | LFQ control | LFQ CG17669 | -log2 ratio | -log10 p value |
| --- | --- | --- | --- | --- | --- |
| Q9VWZ3 | Dhc16F | 1.01829888 | 0.041609539 | 4.613103 | 3.265697 |
| A8Y5B7 | kl-3 | 6.80093659 | 0.430428367 | 3.981888 | 3.182219 |
| A1ZB91 | CG17669 | 1.2915202 | 0.081774953 | 3.981267 | 3.103016 |
| Q5LJP0 | kl-2 | 0.78943234 | 0.224258622 | 1.815652 | 2.822993 |
| Q9VU57 | Cfap43 | 0.55106365 | 0.068451057 | 3.009074 | 2.674799 |
| Q9VJC6 | Dhc36C | 2.82836711 | 0.163285342 | 4.114502 | 2.494444 |
| Q7KVA7 | Dhc62B | 1.00184398 | 0.157084244 | 2.673047 | 2.321502 |
| Q9VZ77 | Dnah3 | 1.08739055 | 0.119223228 | 3.189133 | 2.309416 |
| Q9VKJ5 | Dyx1c1 | 0.05987797 | 0.20546445 | -1.77879 | 2.303495 |
| Q0E8T7 | CG34124 | 1.18667573 | 0.490089781 | 1.275808 | 2.25354 |
| Q5LJN5 | kl-5 | 8.19193092 | 2.814315855 | 1.541419 | 2.208549 |
| Q9VAV5 | Dhc98D | 1.83012501 | 0.712779034 | 1.360415 | 2.116203 |
| Q9VM21 | gudu | 1.62031671 | 0.232234203 | 2.802623 | 2.02667 |
| Q9VVM7 | Wdr92 | 0.27889713 | 0.453183439 | -0.70036 | 1.994956 |
| Q9VJY4 | Dnai2 | 0.67582694 | 0.303152945 | 1.156608 | 1.965192 |
| Q9W0U9 | Dic61B | 0.22810593 | 0.14967333 | 0.607887 | 1.814405 |
| Q9VR52 | Lrrc6 | 0.02411973 | 0.09266751 | -1.94185 | 1.793124 |
| Q9W0F0 | robl62A | 0.25048477 | 0.039268251 | 2.673288 | 1.774009 |
| Q9VCN4 | Ccdc114 | 1.08124333 | 0.208368775 | 2.37548 | 1.746445 |
| Q9VGG6 | Dnali1 | 3.88414271 | 1.440937117 | 1.430589 | 1.638454 |
| Q4V516 | lobo | 0.4429829 | 0.227454095 | 0.961676 | 1.597102 |
| Q9W1V2 | CG3085 | 4.91455102 | 1.939847136 | 1.341117 | 1.576174 |
| Q9VIY1 | CG10750 | 0.03495738 | 0.007402311 | 2.23955 | 1.554517 |
| Q8T3Z0 | Tektin-C | 2.44201942 | 1.224471976 | 0.995915 | 1.457508 |
| Q9W168 | CG16837 | 0.06873574 | 0.028035895 | 1.293785 | 1.453004 |
| Q9W2A3 | CG13501 | 0.13118274 | 0.07513098 | 0.804098 | 1.408852 |

**Table 2.** Dynein heavy chains in *Dnaaf3^ΔCR^* mutant testis proteome

| Protein accession | Gene | Human orthologue | Dynein complex | -log2(ratio) | -log10(p value) |
| --- | --- | --- | --- | --- | --- |
| **ODA** |  |  |  |  |  |
| A8Y5B7 | *kl-3* | DNAH8 | ODA β | 3.981888 | 3.182219 |
| Q5LJN5 | *kl-5* | DNAH17 | ODA γ | 1.541419 | 2.208549 |
| **IDA (single headed)** |  |  |  |  |  |
| Q9VJC6 | *Dhc36C* | DNAH7 | IDA a,b,c | 4.114502 | 2.494444 |
| Q9VZ77 | *Dnah3* | DNAH3 | IDA a,b,c | 3.189133 | 2.309416 |
| Q7KVA7 | *Dhc62B* | DNAH12 | IDA a,b,c | 2.673047 | 2.321502 |
| Q9VWZ3 | *Dhc16F* | DNAH6 | IDA d,e | 4.613103 | 3.265697 |
| **IDA (double headed)** |  |  |  |  |  |
| Q9VAV5 | *Dhc98D* | DNAH10 | IDA f | 1.360415 | 2.116203 |
| Q5LJP0 | *kl-2* | DNAH2 | IDA f | 1.815652 | 2.822993 |

## DISCUSSION

### *Dnaaf3* has conserved functions in ciliary axonemal dynein assembly

Functional analysis establishes that *Drosophila Dnaaf3* (*CG17669*), like human *DNAAF3*, is required for axonemal dynein complex assembly and localisation. Moreover, it is required for dynein localisation in both IFT-dependent motile cilia (Ch neuron dendrites) and IFT-independent flagella (sperm). The *Dnaaf3* phenotype, including the lack of visible ODA and IDA on TEM images, suggests a complete loss of dynein arms from mature cilia/flagella. Together with the cytoplasmic location of Dnaaf3 protein, these features are consistent with a role in the cytoplasmic pre-assembly of dynein complexes, as has been proposed for the human and *Chlamydomonas* orthologues.

In PCD patients with *DNAAF3* mutations, respiratory cilia did not localise several subunits associated with ODA (DNAH5, DNAH9, and DNAI2) or a subunit associated with a subset of single-headed IDAs (DNALI1). *Chlamydomonas pf22* mutant flagella similarly showed loss of subunits associated with ODA and the single-headed IDA forms b and c (see Fig. 7C for summary of dynein forms). However, a subunit of the double-headed form, IDA f (IC140), was still present in mutant flagella (Mitchison et al., 2012) suggesting that this IDA form did not require *DNAAF3* for its assembly. However, in our proteomic analysis we found reduction in abundance of all sperm-expressed HCs in the *Dnaaf3* mutant testes. Notably, this includes the two HCs of double-headed IDA f, both of which were prominently reduced. To reconcile this, we suggest that the presence of IC140 in *Chlamydomonas pf22* mutant flagella does not represent a complete IDA f complex, but just the IC-LC part. Indeed, this IC-LC part is thought to function independently as a motility regulator and has been called the IC138 subcomplex (Bower et al., 2009) or outer-inner dynein linker (OID) (Oda et al., 2013). We suggest, therefore, that the IC138 subcomplex/OID can assemble and localise in the absence of HCs, and moreover its assembly does not require DNAAF3. Interestingly, a parallel to this can be seen in wild-type *Drosophila* Ch neurons: we previously showed that these cells express IC138 subcomplex/OID subunits but not the corresponding IDA f HCs, supporting the presence of an independent IC138 subcomplex/OID in these cilia (zur Lage et al., 2019).

Based on protease sensitivity and antigen presentation in *pf22* mutants, it was hypothesised that abnormal dynein complexes accumulate, leading to the suggestion that PF22 functions at a late step in assembly, possibly in stabilising HCs or in late stage maturation after cytoplasmic pre-assembly (Mitchison et al., 2012). Moreover, cytoplasmic abundances of HCs were not affected, whereas IC2 was increased. This contrasts with other dynein assembly factors, which often show reduction in cytoplasmic levels consistent with protein instability. This was also taken to support the idea that DNAAF3 acts at a different step from other dynein assembly factors. In contrast, in *Drosophila* mutant testes we found reduction in total amounts of many motor components but particularly HCs. We cannot, however, be certain that this reflects cytoplasmic levels in the spermatocytes.

Several DNAAFs have been found to interact during their function, such as DYX1C1/PIH1D3 and ZMYND10/LRRC6 (Moore et al., 2013). In contrast, investigations of protein interactions of DNAAF3 homologues have not so far proved informative (Mitchison et al., 2012). Similarly, we have not been able to identify proteins that interact with *Drosophila* Dnaaf3 by AP-MS (unpublished observations). In the samples, the MS/MS count of Dnaaf3 was large, but few associated proteins were detected, and none appeared interesting from the point of view of dynein biology, protein folding, or transport. The molecular function of DNAAF3 homologues therefore remains mysterious. The inference is that DNAAF3 protein interactions are too transient for detection by immunoprecipitation methods.

### Axonemal dyneins are required for mechanosensation by Ch neurons

By Ca imaging, we show that the Ch neurons of *Dnaaf3* mutant larvae are unable to respond to a vibration stimulus, and the larvae correspondingly do not respond behaviourally to a tone stimulus. This confirms that axonemal dynein motors in Ch neuron cilia are absolutely required for the mechanotransduction process, supporting previous findings in adult auditory Ch neurons, in which mutation of ODA IC or IDA HC subunits caused defects in auditory mechanotransduction (Karak et al., 2015). The molecular role of the axonemal dyneins in mechanotransduction is not fully determined, but they are likely responsible for active mechanical amplification of stimuli as well as adaptation within Ch neuron cilia (Albert and Göpfert, 2015).

### A role for axonemal dynein motors in axoneme stabilisation during IFT-independent flagellogenesis?

Like most cilia, ciliogenesis in *Drosophila* sensory neurons occurs from a basal body docked at the plasma membrane, with the axoneme extending within a bounding membrane. This requires IFT for transport of components within the extending ciliary compartment (i.e. ciliogenesis is compartmentalised). In contrast, *Drosophila* sperm flagellogenesis occurs by axonemal extension within the cytosol in a process that does not require IFT (Basiri et al., 2014; Fingerhut and Yamashita, 2020; Han et al., 2003; Sarpal et al., 2003; Vieillard et al., 2016). Microtubule doublet extension occurs within a membrane ‘ciliary cap’ but the growing axoneme is pushed out into the cytoplasm (Basiri et al., 2014). Once outside the ciliary cap, motors are docked from the cytoplasm (Fingerhut and Yamashita, 2020). Subsequently, sperm undergo a process of individualisation which strips off cytoplasm and surrounds the flagellum axoneme with membrane (Basiri et al., 2014; Fabian and Brill, 2012).

Our analysis suggests that *Drosophila Dnaaf3* is specifically required for axonemal dynein assembly. However, in addition to dynein arm loss, *Dnaaf3* sperm flagella show axoneme disruption with doublets frequently splitting off. Such a phenotype has been noted in all *Drosophila* dynein assembly factor mutants analysed so far (*tilB, Zmynd10, Heatr2, Spag1, Wdr92*) (Diggle et al., 2014; Kavlie et al., 2010; Moore et al., 2013; zur Lage et al., 2018). Moreover, axonemal breakage is seen in mutants of many axonemal dynein subunits themselves (including *gudu, kl-2, kl-3, kl-5, Dhc98D*) (Steinhauer et al., 2019). We suggest that in the absence of a bounding membrane during flagellogenesis, dynein complexes provide a temporary doublet ‘cross-linking’ function to maintain axoneme stability prior to being surrounded by membrane during sperm individualisation. When dynein attachment is compromised, the microtubule doublets will be vulnerable to separation prior to individualisation. This then has a knock-on effect of disrupting individualisation, which also appears compromised in mutants with disrupted axonemes axoneme (Fatima, 2011; Steinhauer et al., 2019).

Since a similar cytosolic mode of assembly is also proposed for the formation of mammalian sperm flagella (Avidor-Reiss and Leroux, 2015), it will be interesting to determine whether defective dynein assembly results in secondary defects in axonemal stability and sperm individualisation of mammalian sperm.

## MATERIALS AND METHODS

### Fly stocks

All fly strains were maintained on standard media at 25°C. The following stock were obtained from the Bloomington Stock Center: *y^1^ w\** P{y^t7.7^=nos-phiC31\int.NLS}X; P{y^t7.7^=CaryP}attP40 (#79604), *y^1^* M{RFP3xP3.PB]GFP^E.3xP3^=vas-Cas9}ZH-2A (#52669), *w^1118^*/FM7c (#51323), *w^1118^* (#3605), *w\** P{w^+mC^=iav-Gal4.K}3 (#52273), *w\** P{y^+t7.7^ w^+mC^=5xUAS-GCaMP6f}attP2 (#91989), and P{w^+mC^=UAS-Dcr-2.D}1 (#24648). The w; Tft/CyO; Bam-VP16-Gal4 was a gift from H. White-Cooper and the sca-Gal4 line from M. Mlodzik. The lines for the RNAi experiments, GD control (ID 60000) and CG17669 RNAi (ID GD36538) were obtained from the Vienna *Drosophila* Resource Center (VDRC) (Dietzl et al., 2007).

### *In situ* hybridisation

*In situ* hybridisation on overnight wild-type embryos was carried out according to (zur Lage et al., 2019) and on 3-4-day old adult testes according to (Morris et al., 2009). The mounted slides were photographed on an Olympus AX70 upright microscope. The RNA probe for *Dnaaf3* was generated by T7 RNA polymerase *in vitro* transcription from a gene fragment isolated by PCR using the primers CAGACTGGACCGTTTTGTGG and GTAATACGACTCACTATAGGGCCCACATGTTCTTGCCGTTGA. The latter includes the T7 RNA polymerase recognition sequence.

### mVenus fusion gene construction

The CG17669::mVenus fusion construct was designed to include the CG17669 ORF and upstream regulatory sequences, with mVenus fused in-frame C-terminally. PCR primers used to amplify the CG17669 sequence were GGGGACAAGTTTGTACAAAAAAGCAGGCTAGTGTTGCATACGAAAGGGT and GGGGACCACTTTGTACAAGAAAGCTGGGTCTGGGAAGTTAGTTTTTGGTT. The cloning was performed according to the Gateway two-step recombination system (Life Technologies). After the BP reaction, the construct was transferred into the pBID-UASC-GV vector (Wang et al., 2012) via the LR reaction in order to generate the expression clone pBID-UASC-CG17669::mVenus. The construct was subsequently microinjected into the attp40 line.

### CRISPR/Cas9-guide RNA expression construction

A *CG17669 CRISPR/Cas9* mutant was constructed by a mini-white gene substitution as described by (zur Lage et al., 2018). Two pairs of sense and antisense oligonucleotides (oligos) were designed for guide RNAs (gRNA I and II, respectively) using the CRISPR optimal target finder (http://tools.flycrispr.molbio.wisc.edu/targetFinder/index.php) based on the *CG17669* sequence. Guides were, gRNA I: CTTCGGAGGATGAGCATGCACGCG and AAACCGCGTGCATGCTCATCCTCC; gRNA II: CTTCGAACCTGCTGAAGAGTATAC and AAACGTATACTCTTCAGCAGGTTC. The guides were phosphorylated and annealed before cloning into the *BbsI* restriction site of pBFv-U6.2 and pBFv-U6.2B (Kondo and Ueda, 2013), respectively. For both guides to be expressed from the same plasmid, an *EcoRI-NotI* fragment from pBFv-U6.2 containing the gRNA I sequence was subsequently cloned into the gRNA II-containing plasmid pUBFv-U6.2B. For homology-directed repair, homology arms were amplified from genomic DNA using the following primer pairs: left arm – GAT<u>GCATGC</u>CGTCAAACAACAGCCAAAAG and GAT<u>GGTACC</u>ACTTCCACTTCCACCCTGGT; right arm – GGG<u>AGATCT</u>TACTGGTCTAGTAATTGAAG and CAA<u>CTCGAGA</u>TCCATAATTGCTGGCAACC. The left homology arm was cloned into the *SphI* and *Asp*718I restriction sites (underlined) of the 5’ multiple cloning site of pRK2 (Huang et al., 2008), whereas the right homology arm was cloned into the *BglII* and *XhoI* sites of the 3’ multiple cloning site of the same vector. The guide RNA and the homology arm constructs were injected into the vas-Cas9 line. Substitution of the *CG17669* ORF by mini-white (the *Dnaaf3^ΔCR^* allele) was confirmed by sequencing.

### Immunofluorescence

Embryo and pupal antenna immunohistochemistry was carried out according to (zur Lage et al., 2018). The fixing and staining of *Drosophila* testis was described in Sitaram et al. 2014 (Sitaram et al., 2014). The following primary antibodies were used: rabbit anti-GFP antibody (1:500, Life Technologies, A11122), mouse anti-Futsch antibody (1:200, Developmental Studies Hybridoma Bank, 22C10-s), mouse anti-acetylated tubulin (1:1000, Sigma, T6793) and rabbit anti-Dnah5 antibody (1:2000, see below) and the secondary antibodies were goat anti-Rabbit antibody (1:500, Alexa Fluor 488, Life Technologies, A11008) and goat anti-Mouse antibody (1:500, Alexa Fluor 568, Life Technologies, A11019). DNA in adult testes was stained with To-Pro-3 (1:1000, Life Technologies, T3605) solution in the dark for 15min. After several washes, the samples were mounted on slides with 85% glycerol and 2.5% propyl gallate (Sigma, P3130). The slides were imaged using a Zeiss LSM-5 PASCAL/Axioskop 2 confocal microscope and processed with Fiji.

### Dnah5 (CG9492) antibody generation

A 744-bp sequence was amplified from exon 5 of the *Drosophila* CG9492 gene and cloned into the *Eco*RI restriction site (underlined) of the expression vector pRSET-A using the following primers: left GGG<u>GAATTC</u>TATATTGCGGAGTGATCGATG and right GAG<u>GAATTC</u>TGACTTGACCATCAGCGCGGATG. The protein was expressed and the antirabbit antibody raised and purified by Proteintech.

### Fertility assay

Single males were mated with two virgin female Oregon-R flies. The flies were allowed to mate at 25°C for 48 h before being transferred to a second batch of fresh vials. After another 48 h, the female flies were removed, and males were dissected to examine the testes for sperm motility. During the period of 10-19 days after the removal of the flies, the progeny in each vial of the second batch was counted.

### Climbing assay

A group of 10 mated 3-4 day old female flies were placed in a cylinder divided vertically into four 5-cm quadrants. After a 1-min recovery period, they were banged to the bottom of the cylinder, and then the height climbed in 10 s was recorded. The quadrant at the top is given a weight of 4 followed by 3, 2 and 1 to the quadrant at the bottom, and the Climbing Index = Σ (the numbers of flies in each quadrant) x (each quadrant weight) / the total number of flies. The assay was repeated for 5 groups of flies.

### Larval hearing assay

Groups of 5 third instar larvae were placed on a grape juice agar plate positioned on a speaker. After 15 s, a 1-s tone of 1000 Hz was delivered by the speaker. For each larva, body contraction was scored (yes=1; no=0) during a 1-s interval prior to the tone and during the tone. Each larva was tested three times (at 30-s intervals). The Response Index for test was the sum of the larval scores (0–5) and these were then averaged for the batch. Mean Response Index was determined from 3 trials (15 larvae total) for each genotype. Significance of differences in Mean Response Index prior and during the tone for each genotype were analysed by 2-way ANOVA followed by Sidak’s test for multiple comparisons.

### Calcium imaging of larval Ch neurons

Third instar larvae with Ch neuron-specific Gal4 (iav-Gal4) driving GCaMP (iav-Gal4/UAS-GCaMP6f) were filleted in Ca^2+^-free Ringer solution (140 mM NaCl, 2 mM KCl, 6 mM MgCl_2_, 5 mM Hepes-NaOH, 36 mM sucrose, pH 7.2). After dissection and washing, saline was replaced with Ringer solution (130 mM NaCl, 5 mM KCl, 2 mM MgCl_2_, 2 mM CaCl_2_, 5 mM Hepes-NaOH, 36 mM sucrose, pH 7.2). The fillet prep was imaged using a Zeiss AX10 Examiner A1 fluorescence microscope with a 40x water immersion objective. Imaging was achieved by a Q-imaging WLS LED illumination unit and Photometrics Prime sCMOS camera. Video was recorded at 20 fps for 40 s. Response was recorded as peak ΔF/F_0_ (%) of mean fluorescence in a Region of Interest (ROI) bounding the paired axon terminals of the Ch neurons in the ventral nerve cord (VNC) when stimulated by 1024-Hz tuning fork (52.4 ± 2.8dB, n = 10 tones). For each fillet prep, three stimuli were presented at 10 s intervals. Data were analysed in FIJI and Microsoft Excel. F_0_ for ΔF/F_0_ (%) was defined as mean F of: (0s to onset of 1^st^ peak) + (end of 1^st^ peak to onset of 2^nd^ peak) + (end of 2^nd^ peak to onset of 3^rd^ peak). The ‘peaks’ used in calculation for mean peak of *Dnaaf3^ΔCR^* homozygotes were derived from mean time(s) of recording that correspond to peaks in *Dnaaf3^ΔCR^* heterozygotes, to allow comparison despite lack of peaks present in the homozygote. Mean response to 1024-Hz stimulation was generated by aligning all peaks in recordings (from 0.5 s before onset of peak to end of peak + 0.5 s) where F_0_ was defined as mean F recorded 0.5 s before onset of peak.

### Transmission electron microscopy (TEM)

Whole heads (proboscis removed to facilitate infiltration) or testes of newly eclosed adult males were removed and rinsed in 0.1M phosphate buffer (pH7.4). The samples were fixed in freshly made 2.5% glutaraldehyde, 2% paraformaldehyde in 0.1M sodium phosphate buffer (pH7.4) solution overnight at 4°C. Post fixing, the samples were first rinsed four times and then washed three times for 20min in 0.1M phosphate buffer at room temperature. The head or testes samples were post-fixed and imaged by Tracey Davey at the Electron Microscopy Research Services, Newcastle University Medical School, using a Philips CM100 CompuStage (FEI) microscope and an AMT CCD camera.

### Protein expression analysis of testes by MS

Four replicates of 30 pairs of 3-4 day old testes were dissected for the *CG17669* CRISPR homozygous mutant line and the Cas9 injection line (control). Sample processing and analysis of the label-free mass-spectrometry was carried out as described by (zur Lage et al., 2018). Proteins with average LFQ value <0.05 in control testes were excluded from further analysis. The mass spectrometry proteomics data have been deposited in the ProteomeXchange Consortium via the PRIDE (Perez-Riverol et al., 2019) partner repository with the dataset identifier PXD025409.

## ACKNOWLEDGEMENTS

We thank Olaf Tomola for preliminary analysis of the proteomics dataset. We are very grateful to Dr Barry Denholm (University of Edinburgh) for use of his Ca-imaging microscope.

## REFERENCES

Albert, J. T. and Göpfert, M. C. (2015). Hearing in Drosophila. Curr. Opin. Neurobiol. 34, 79–85.

Avidor-Reiss, T. and Leroux, M. R. (2015). Shared and Distinct Mechanisms of Compartmentalized and Cytosolic Ciliogenesis. Curr. Biol. 25, R1143–R1150.

Basiri, M. L., Ha, A., Chadha, A., Clark, N. M., Polyanovsky, A., Cook, B. and Avidor-Reiss, T. (2014). A migrating ciliary gate compartmentalizes the site of axoneme assembly in drosophila spermatids. Curr. Biol. 24, 2622–2631.

Bower, R., Vanderwaal, K., O ‘toole, E., Fox, L., Perrone, C., Mueller, J., Wirschell, M., Kamiya, R., Sale, W. S. and Porter, M. E. (2009). IC138 Defines a Subdomain at the Base of the I1 Dynein That Regulates Microtubule Sliding and Flagellar Motility. Mol. Biol. Cell 20, 3055–3063.

Dietzl, G., Chen, D., Schnorrer, F., Su, K., Barinova, Y., Fellner, M., Gasser, B., Kinsey, K., Oppel, S., Scheiblauer, S., et al.(2007). A genome-wide transgenic RNAi library for conditional gene inactivation in Drosophila. Nature 448, 151–156.

Diggle, C. P., Moore, D. J., Mali, G., zur Lage, P., Ait-Lounis, A., Schmidts, M., Shoemark, A., Garcia Munoz, A., Halachev, M. R., Gautier, P., et al.(2014). HEATR2 Plays a Conserved Role in Assembly of the Ciliary Motile Apparatus. PLoS Genet. 10, e1004577.

Fabian, L. and Brill, J. A. (2012). Drosophila spermiogenesis. Spermatogenesis 2, 197–212.

Fatima, R. (2011). Drosophila Dynein Intermediate Chain Gene, Dic61B, Is Required for Spermatogenesis. PLoS One 6, e27822.

Fingerhut, J. and Yamashita, Y. (2020). mRNA localization mediates maturation of cytoplasmic cilia in Drosophila spermatogenesis. J. Cell Biol. 219, e202003084.

Fok, A. K., Wang, H., Katayama, A., Aihara, M. S. and Allen, R. D. (1994). 22S axonemal dynein is preassembled and functional prior to being transported to and attached on the axonemes. Cell Motil. Cytoskeleton 29, 215–224.

Fowkes, M. E. and Mitchell, D. R. (1998). The role of preassembled cytoplasmic complexes in assembly of flagellar dynein subunits. Mol. Biol. Cell 9, 2337–47.

Guo, Z., Chen, W., Huang, J., Wang, L. and Qian, L. (2019). Clinical and genetic analysis of patients with primary ciliary dyskinesia caused by novel DNAAF3 mutations. J. Hum. Genet. 64, 711–719.

Han, Y. G., Kwok, B. H. and Kernan, M. J. (2003). Intraflagellar Transport Is Required in Drosophila to Differentiate Sensory Cilia but Not Sperm. Curr. Biol. 13, 1679–1686.

Horani, A. and Ferkol, T. W. (2013). Molecular Genetics of Primary Ciliary Dyskinesia. In eLS, p. Chichester, UK: John Wiley & Sons, Ltd.

Kakihara, Y. and Houry, W. A. (2012). The R2TP complex: discovery and functions. Biochim. Biophys. Acta 1823, 101–7.

Karak, S., Jacobs, J. S., Kittelmann, M., Spalthoff, C., Katana, R., Sivan-Loukianova, E., Schon, M. A., Kernan, M. J., Eberl, D. F. and Göpfert, M. C. (2015). Diverse Roles of Axonemal Dyneins in Drosophila Auditory Neuron Function and Mechanical Amplification in Hearing. Sci. Rep. 5, 17085.

Kavlie, R. G., Kernan, M. J. and Eberl, D. F. (2010). Hearing in Drosophila Requires TilB, a Conserved Protein Associated With Ciliary Motility. Genetics 185, 177–188.

King, S. M. (2016). Axonemal Dynein Arms. Cold Spring Harb. Perspect. Biol. 8, a028100.

Knowles, M. R., Ostrowski, L. E., Loges, N. T., Hurd, T., Leigh, M. W., Huang, L., Wolf, W. E., Carson, J. L., Hazucha, M. J., Yin, W., et al.(2013). Mutations in SPAG1 cause primary ciliary dyskinesia associated with defective outer and inner dynein arms. Am. J. Hum. Genet. 93, 711–720.

Kondo, S. and Ueda, R. (2013). Highly Improved Gene Targeting by Germline-Specific Cas9 Expression in Drosophila. Genetics 195, 715–721.

Laurençon, A., Dubruille, R., Efimenko, E., Grenier, G., Bissett, R., Cortier, E., Rolland, V., Swoboda, P. and Durand, B. (2007). Identification of novel regulatory factor X (RFX) target genes by comparative genomics in Drosophila species. Genome Biol. 8, R195.

Lin, J., Yin, W., Smith, M., Song, K., Leigh, M., Zariwala, M., Knowles, M., Ostrowski, L. and Nicastro, D. (2014). Cryo-electron tomography reveals ciliary defects underlying human RSPH1 primary ciliary dyskinesia. Nat. Commun. 53, 160.

Mali, G., Yeyati, P., Mizuno, S., Keighren, M., zur Lage, P., Garcia-Munoz, A., Shimada, A., Takeda, H., Edlich, F., Takahashi, S., et al.(2018). ZMYND10 functions in a chaperone relay during axonemal dynein assembly. Elife 7, e34389.

Mitchison, H. M. and Valente, E. M. (2017). Motile and non-motile cilia in human pathology: from function to phenotypes. J. Pathol. 241, 294–309.

Mitchison, H. M., Schmidts, M., Loges, N. T., Freshour, J., Dritsoula, A., Hirst, R. A., O’Callaghan, C., Blau, H., Al Dabbagh, M., Olbrich, H., et al.(2012). Mutations in axonemal dynein assembly factor DNAAF3 cause primary ciliary dyskinesia. Nat. Genet. 44, 381–389.

Moore, D. J., Onoufriadis, A., Shoemark, A., Simpson, M. A., Zur Lage, P. I., De Castro, S. C., Bartoloni, L., Gallone, G., Petridi, S., Woollard, W. J., et al.(2013). Mutations in ZMYND10, a gene essential for proper axonemal assembly of inner and outer dynein arms in humans and flies, cause primary ciliary dyskinesia. Am. J. Hum. Genet. 93, 346–356.

Morris, C., Benson, E. and White-Cooper, H. (2009). Determination of gene expression patterns using in situ hybridization to Drosophila testes. Nat. Protoc. 4, 1807–1819.

Newton, F. G., zur Lage, P. I., Karak, S., Moore, D. J., Göpfert, M. C. and Jarman, A. P. (2012). Forkhead Transcription Factor Fd3F Cooperates with Rfx to Regulate a Gene Expression Program for Mechanosensory Cilia Specialization. Dev. Cell 22, 1221–1233.

Oda, T., Yagi, T., Yanagisawa, H. and Kikkawa, M. (2013). Identification of the outer-inner dynein linker as a hub controller for axonemal dynein activities. Curr. Biol. 23, 656–664.

Omran, H., Kobayashi, D., Olbrich, H., Tsukahara, T., Loges, N. T., Hagiwara, H., Zhang, Q., Leblond, G., O’Toole, E., Hara, C., et al.(2008). Ktu/PF13 is required for cytoplasmic pre-assembly of axonemal dyneins. Nature 456, 611–6.

Paff, T., Loges, N. T., Aprea, I., Wu, K., Bakey, Z., Haarman, E. G., Daniels, J. M. A., Sistermans, E. A., Bogunovic, N., Dougherty, G. W., et al.(2017). Mutations in PIH1D3 Cause X-Linked Primary Ciliary Dyskinesia with Outer and Inner Dynein Arm Defects. Am. J. Hum. Genet. 100, 160–168.

Perez-Riverol, Y., Csordas, A., Bai, J., Bernal-Llinares, M., Hewapathirana, S., Kundu, D. J., Inuganti, A., Griss, J., Mayer, G., Eisenacher, M., et al.(2019). The PRIDE database and related tools and resources in 2019: Improving support for quantification data. Nucleic Acids Res. 47, D442–D450.

Porter, M. E. (2017). Ciliary and flagellar motility and the nexin-dynein regulatory complex. In Dyneins: The Biology of Dynein Motors: Second Edition, pp. 299–335.

Sarpal, R., Todi, S. V., Sivan-Loukianova, E., Shirolikar, S., Subramanian, N., Raff, E. C., Erickson, J. W., Ray, K. and Eberl, D. F. (2003). Drosophila KAP Interacts with the Kinesin II Motor Subunit KLP64D to Assemble Chordotonal Sensory Cilia, but Not Sperm Tails. Curr. Biol. 13, 1687–1696.

Sitaram, P., Hainline, S. G. and Lee, L. A. (2014). Cytological Analysis of Spermatogenesis: Live and Fixed Preparations of <em>Drosophila</em> Testes. J. Vis. Exp. 83,.

Steinhauer, J., Statman, B., Fagan, J. K., Borck, J., Surabhi, S., Yarikipati, P., Edelman, D. and Jenny, A. (2019). Combover interacts with the axonemal component Rsp3 and is required for Drosophila sperm individualization. Dev. 146, dev179275.

Tarkar, A., Loges, N. T., Slagle, C. E., Francis, R., Dougherty, G. W., Tamayo, J. V, Shook, B., Cantino, M., Schwartz, D., Jahnke, C., et al.(2013). DYX1C1 is required for axonemal dynein assembly and ciliary motility. Nat. Genet. 45, 995–1003.

Vaughan, C. K. (2014). Hsp90 Picks PIKKs via R2TP and Tel2. Structure 22, 799–800.

Vieillard, J., Paschaki, M., Duteyrat, J. L., Augière, C., Cortier, E., Lapart, J. A., Thomas, J. and Durand, B. (2016). Transition zone assembly and its contribution to axoneme formation in Drosophila male germ cells. J. Cell Biol. 214, 875–889.

Wang, J.-W., Beck, E. S. and McCabe, B. D. (2012). A Modular Toolset for Recombination Transgenesis and Neurogenetic Analysis of Drosophila. PLoS One 7, e42102.

zur Lage, P., Stefanopoulou, P., Styczynska-Soczka, K., Quinn, N., Mali, G., von Kriegsheim, A., Mill, P. and Jarman, A. P. (2018). Ciliary dynein motor preassembly is regulated by Wdr92 in association with HSP90 co-chaperone, R2TP. J. Cell Biol. 217, 2583–2598.

zur Lage, P., Newton, F. G. and Jarman, A. P. (2019). Survey of the Ciliary Motility Machinery of Drosophila Sperm and Ciliated Mechanosensory Neurons Reveals Unexpected Cell-Type Specific Variations: A Model for Motile Ciliopathies. Front. Genet. 10,.

