## Supplemental figures, table, and movie captions for "The *Drosophila* orthologue of the primary ciliary dyskinesia-associated gene, *DNAAF3*, is required for axonemal dynein assembly"

**Fig. S1**

*Dnaaf3-mVenus* fusion gene was designed to include the X box (consensus nucleotide sequence RYYRYYN(1–3)RRNRAC) (Laurençon et al., 2007) and Forkhead binding motif (F motif) (consensus nucleotide sequence RYMAAYA) (Newton et al., 2012) motifs in the upstream regulatory region to keep the spatial-temporal pattern of the expression from its own promoter.

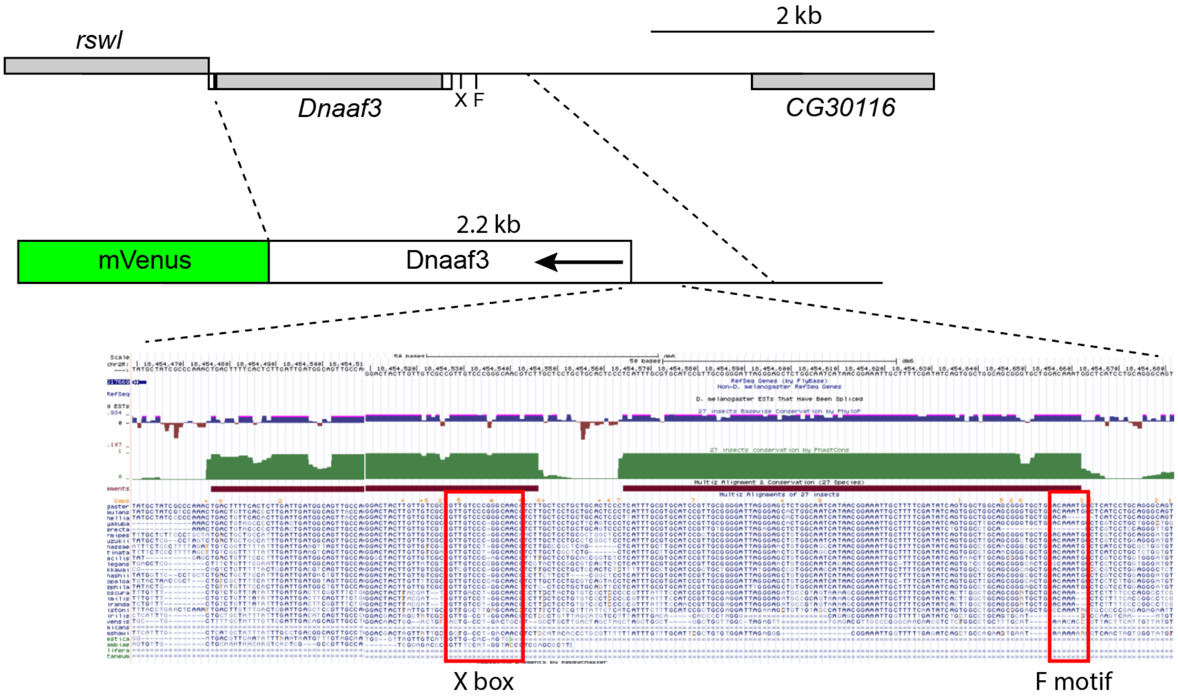

**Fig. S2**

*Dnaaf3* mRNA expression (A) Adult testis. The large nuclei identify the expressing cells as spermatocytes. No expression is seen in mature sperm. (B) Stage 16 embryo. Expression is observed in the segmentally repeated locations expected for larval Ch neurons (see Fig. 1A).

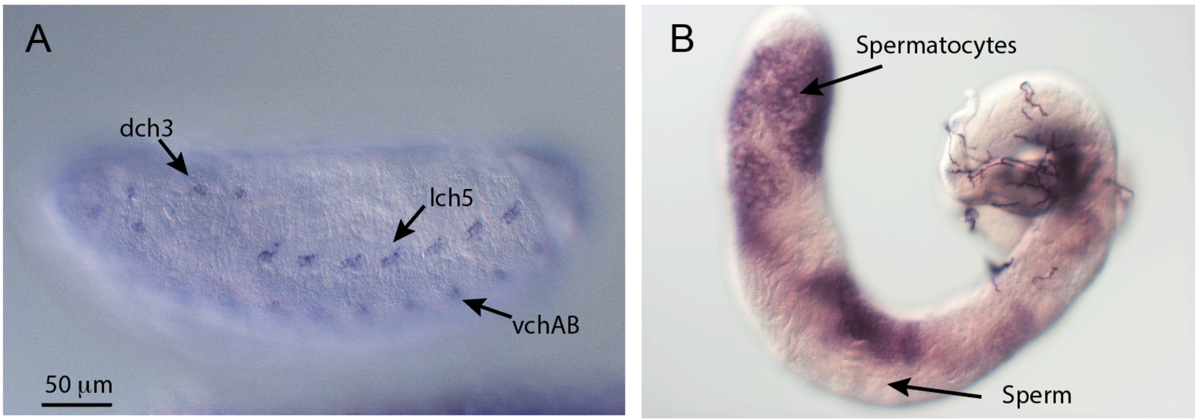

**Movie 1:** Sperm from *Bam*Gal4 fly (control for RNAi)

**Movie 2:** Sperm from *Bam*Gal4; UAS-*Dnaaf3* RNAi fly (*Dnaaf3* knockdown)

**Movie 3:** Sperm from *Dnaaf3*$^{\Delta CR}$ homozygote

**Movie 4:** Sperm from *Dnaaf3*$^{\Delta CR}$ homozygote with *Dnaaf3-mVenus* fusion gene (mutant rescue)

### Table S1. Ciliary motility proteins detected in adult testes, sorted by abundance

| Majority protein IDs | Gene names | Average LFQ control* | Average LFQ CG17669* | -log2 ratio | -log10 p value |
|---|---|---|---|---|---|
| Q9VH07 | pont | 9.87783848 | 9.62517004 | 0.037383 | 0.032098 |
| Q8INT5 | Dnaaf1 | 9.14370354 | 11.50210626 | -0.33105 | 0.265354 |
| Q5LJN5 | kl-5 | 8.19193092 | 2.814315855 | 1.541419 | 2.208549 |
| Q9V3K3 | rept | 7.88284253 | 8.65178034 | -0.13428 | 0.235542 |
| Q9V3M9 | Tektin-A | 7.25402673 | 2.979805372 | 1.283564 | 1.186591 |
| A8Y5B7 | kl-3 | 6.80093659 | 0.430428367 | 3.981888 | 3.182219 |
| Q9VBA1 | Spag1 | 6.39372796 | 6.470997003 | -0.01733 | 0.011752 |
| Q9VU41 | Zmynd10 | 5.59104156 | 7.163702465 | -0.35759 | 0.363388 |
| Q9W1V2 | CG3085 | 4.91455102 | 1.939847136 | 1.341117 | 1.576174 |
| Q9VGG6 | Dnali1 | 3.88414271 | 1.440937117 | 1.430589 | 1.638454 |
| Q9VGC1 | CG10014 | 3.6990868 | 2.098445308 | 0.817848 | 0.688281 |
| Q9V3E9 | Rpap3 | 3.05702371 | 3.072928769 | -0.00749 | 0.0058 |
| Q9W1D3 | Rsph4a | 3.04296406 | 2.084000363 | 0.546122 | 0.926048 |
| Q9VJC6 | Dhc36C | 2.82836711 | 0.163285342 | 4.114502 | 2.494444 |
| Q8T3Z0 | Tektin-C | 2.44201942 | 1.224471976 | 0.995915 | 1.457508 |
| Q8INF7 | Heatr2 | 2.35414183 | 2.489974686 | -0.08093 | 0.169468 |
| Q9VT21 | CG8336 | 2.33143917 | 2.505177094 | -0.10369 | 0.116157 |
| Q9VAV5 | Dhc98D | 1.83012501 | 0.712779034 | 1.360415 | 2.116203 |
| Q9VM21 | gudu | 1.62031671 | 0.232234203 | 2.802623 | 2.02667 |
| Q9VUG3 | Pih1D3 | 1.57346805 | 1.741292636 | -0.14621 | 0.1014 |
| Q9VZH1 | CG18675 | 1.47753788 | 1.76610829 | -0.25738 | 0.309647 |
| Q9VN57 | CG17387 | 1.41627716 | 1.324956294 | 0.096159 | 0.070752 |
| Q24117 | ctp | 1.36907896 | 1.078564964 | 0.344093 | 0.207812 |
| Q9VK29 | Rsph1 | 1.35813245 | 0.991262186 | 0.454286 | 0.621901 |
| **A1ZB91** | **CG17669** | **1.2915202** | **0.081774953** | **3.981267** | **3.103016** |
| Q0E8T7 | CG34124 | 1.18667573 | 0.490089781 | 1.275808 | 2.25354 |
| Q8T415 | Centrin | 1.16685798 | 0.984236486 | 0.245552 | 0.207075 |
| Q9VYR5 | CG18130 | 1.10884709 | 0.581556947 | 0.931068 | 0.909879 |
| Q9VZ77 | Dnah3 | 1.08739055 | 0.119223228 | 3.189133 | 2.309416 |
| Q9VCN4 | Ccdc114 | 1.08124333 | 0.208368775 | 2.37548 | 1.746445 |

| | | | | | |
|---|---|---|---|---|---|
| Q94524 | Dlc90F | 1.08013754 | 0.998358295 | 0.113585 | 0.088463 |
| Q8MT08 | Gas8 | 1.05840147 | 0.268884548 | 1.976828 | 1.168448 |
| Q9VWZ3 | Dhc16F | 1.01829888 | 0.041609539 | 4.613103 | 3.265697 |
| Q7KVA7 | Dhc62B | 1.00184398 | 0.157084244 | 2.673047 | 2.321502 |
| Q9VRY7 | CG10099 | 0.91960338 | 0.606549278 | 0.600387 | 0.664016 |
| Q8T3V7 | Rsph9 | 0.86288384 | 0.617610339 | 0.482469 | 0.659223 |
| Q9VC34 | CG6980 | 0.83779304 | 1.043301876 | -0.31649 | 0.301679 |
| Q5LJP0 | kl-2 | 0.78943234 | 0.224258622 | 1.815652 | 2.822993 |
| A0A0B7P7M6 | l(2)41Ab | 0.76685658 | 0.609264688 | 0.331888 | 0.30116 |
| Q9VS90 | Wdr63 | 0.76016435 | 0.510863279 | 0.573374 | 1.099499 |
| Q9VJY4 | Dnai2 | 0.67582694 | 0.303152945 | 1.156608 | 1.965192 |
| Q9VA28 | CG15547 | 0.55961706 | 0.304514523 | 0.877929 | 0.757275 |
| Q9VU57 | Cfap43 | 0.55106365 | 0.068451057 | 3.009074 | 2.674799 |
| Q0E9G3 | Dnaaf2 | 0.47060279 | 0.581374511 | -0.30496 | 0.22944 |
| Q4V516 | lobo | 0.4429829 | 0.227454095 | 0.961676 | 1.597102 |
| Q9W3J8 | CG10958 | 0.3214128 | 0.131678316 | 1.28741 | 1.370424 |
| Q9VVM7 | Wdr92 | 0.27889713 | 0.453183439 | -0.70036 | 1.994956 |
| Q9W0F0 | robl62A | 0.25048477 | 0.039268251 | 2.673288 | 1.774009 |
| Q4QPW2 | CG10822 | 0.23635937 | 0.124063924 | 0.929898 | 0.591768 |
| Q9W0U9 | Dic61B | 0.22810593 | 0.14967333 | 0.607887 | 1.814405 |
| A1Z8T9 | CG8407 | 0.21119504 | 0.090270929 | 1.226243 | 0.634525 |
| Q7KMS3 | robl | 0.17293244 | 0.22301487 | -0.36693 | 0.193481 |
| Q0KI05 | Dnah9 | 0.15915179 | 0.020345612 | 2.967614 | 0.376532 |
| Q9W2A3 | CG13501 | 0.13118274 | 0.07513098 | 0.804098 | 1.408852 |
| Q9W168 | CG16837 | 0.06873574 | 0.028035895 | 1.293785 | 1.453004 |
| Q9VKJ5 | Dyx1c1 | 0.05987797 | 0.20546445 | -1.77879 | 2.303495 |
| Q9V9P1 | CG10834 | 0.04958531 | *5.00819E-10* | | |
| Q7K4X4 | Cfap53 | 0.04266028 | 0.11420404 | -1.42065 | 0.929194 |
| Q9VIY1 | CG10750 | 0.03495738 | 0.007402311 | 2.23955 | 1.554517 |
| Q9VK58 | Pih1D1 | 0.03250981 | 0.03428731 | -0.0768 | 0.097999 |
| Q9W3L0 | Dnai2 | 0.02907692 | 0.004135815 | 2.813631 | 0.631246 |
| Q9VQA6 | robl22E | 0.02688946 | 0.036787338 | -0.45217 | 0.136057 |
| Q9VR52 | Lrrc6 | 0.02411973 | 0.09266751 | -1.94185 | 1.793124 |

| | | | | | |
|---|---|---|---|---|---|
| Q7KUM9 | CG7276 | 0.01382293 | 0.005903806 | 1.227346 | 0.44837 |
| A1Z8V5 | Iqcd | 0.01227507 | 0.001742399 | 2.816585 | 1.135211 |
| A0A0B4LH20 | CG9492 | 0.00647783 | 0.006053899 | 0.097646 | 0.016186 |
| Q4V5H1 | CG17266 | 0.0048374 | 0.007403708 | -0.61402 | 0.278398 |
| Q9VJ18 | robl37BC | 0.00099315 | *5.00819E-10* | | |
| P47947 | TpnC41C | *5.7109E-10* | *5.00819E-10* | | |
| A1Z8J9 | CG13202 | *5.7109E-10* | 0.017887853 | | |
| A8JUM9 | Casc1 | *5.7109E-10* | *5.00819E-10* | | |
| Q9VJC5 | Maats1 | *5.7109E-10* | *5.00819E-10* | | |

* italics represent imputed values. Proteins in red were excluded from further analysis due to very low detection level (average LFQ in control <0.05).
